# *Escherichia coli* transcription factors RapA and SspA play opposing roles in tolerance to replication/ transcription conflicts after DNA damage

**DOI:** 10.64898/2026.01.23.701399

**Authors:** Thalia H. Sass, Elena Delicado Dominguez, Susan T. Lovett

## Abstract

The ability to clear transcription complexes from DNA is especially important after DNA damage that produces replication stress. Using the bacterium *E. coli,* we show here that mutations in RNA polymerase that reduce termination, inhibitors of Rho-dependent termination, and inversion of a highly transcribed ribosomal RNA operon both enhance sensitivity to the quinolone ciprofloxacin (CPX); and identify two transcription factors, SspA and RapA, that impact these effects in opposite ways.We demonstrate that the *rapA* promoter is induced by CPX, independent of the LexA/RecA SOS response but is dependent on DnaA. Previous work has shown that RapA is expressed highest in rapidly growing cells whereas SspA levels respond to starvation.The factors have opposing effects on tolerance to chronic exposure to CPX, with RapA promoting cell growth and SspA inhibiting it. Functional SspA is also required for the CPX toxicity of the rRNA operon inversion; in *sspAΔ* mutants it has no negative consequence. In otherwise wild-type cells, loss of RapA has little effect except in strains lacking RNase HI, the enzyme that removes RNA/DNA hybrids from DNA. However, in cells lacking SspA, RapA strongly promotes survival, suggesting that SspA must block positive effects of RapA on tolerance. The RapA requirement for CPX tolerance is not relieved by RNase HI overexpression and therefore RapA must be not be merely preventing R-loop formation. RapA also in some way promotes the use of RNA loops to initiate DNA replication in the absence of DnaA. We propose that SspA stabilizes stalled or post-termination RNAP/DNA complexes and that the presence of SspA prevents RapA release of these complexes.

## INTRODUCTION

DNA replication occurs on DNA that is actively transcribed therefore, the conflicts between replication and transcription machinery is a fundamental problem encountered in cells of all organisms. In the bacterium *E. coli*, the DNA polymerase III replisome is about 20 times faster than transcription machinery, making collisions inevitable (Mirkin and Mirkin 2005). It is estimated that the *E. coli* K12 chromosome has 4000 actively transcribing RNA polymerase molecules at any given time during fast growth (doubling time of 20 minutes), as would occur with growth in rich culture medium such as LB (Bremer and Dennis 2008). Thus, a single round of replication requires resolution of multiple collision events with transcription elongation complexes (TEC). Most of these active transcription complexes, about 90% during fast growth, are found at the 7 operons that encode ribosomal RNA (rRNA) (Bremer and Dennis 2008).

Collision of the replisome with TECs can be in two orientations: head-on or co-directional, with the former more problematic, as evident from in vivo studies from a wide variety of organisms (French 1992, Srivatsan,Tehranchi et al. 2010, Cooper, Harada et al. 2021, Boubakri, de Septenville et al. 2010). *E. coli* possesses 7 operons devoted to the synthesis of ribosomal RNA; not surprisingly, all 7 are co-oriented with the direction of replication from its single origin of replication, *oriC,* thereby ensuring that collisions between replication and transcription machinery at these heavily transcribed loci would be the more benign, co-directional, type. Replication/transcription conflicts can also lead to the formation of stable RNA/DNA hybrid molecules, known as “R-loops” (Lang et al. 2017), which can be deleterious.

Problems associated with collisions of transcription complexes may be exacerbated by DNA damage and other sources of replication problems. An inversion of a highly transcribed rRNA operon such that it produces a head-on collision with the replisome is not normally problematic but becomes more lethal when combined with mutants the DNA polymerase III replication clamp loader protein HolC. Mutants in RNA polymerase beta subunit that increase processivity and discourage TEC dissociation are deleterious in *holCΔ*; conversely mutants that promote stalling and termination suppress *holCΔ* inviability (Cooper et al. 2021). Under replication stress conditions promoted by nalidixic acid or by loss of *holC*, Rho-mediated transcriptional termination becomes essential for tolerance and growth (Washburn and Gottesman 2011; Myka and Gottesman 2019; Myka et al. 2019; Cooper et al. 2021).

Our recent work suggested a potential role for the RapA transcription factor in tolerance of replication stress. Our RNA-Seq studies of gene expression after replication stress showed that the transcription recycling factor, RapA was strongly induced by replication inhibitors azidothymidine (AZT) (Sass and Lovett 2024) and ciprofloxacin (CPX) (unpublished data) in a manner independent of the SOS DNA damage response controlled by LexA and RecA. Induction of *rapA* was noted previously for UV (Courcelle et al. 2001) and for norfloxacin in microarray studies (Dwyer et al. 2007). RapA is an RNA polymerase-associated SWI2/SNF2 family protein (initially named “HepA”) that promotes RNA polymerase (RNAP) recycling during in vitro transcription (Sukhodolets et al. 2001). RapA associates with the RNAP core in a way that is proposed to modulate the RNAP clamp (Qayyum et al. 2021; Brewer et al. 2025) that affects the interaction with nucleic acids. RapA is able to recycle an RNA-free RNAP post-termination complex and modulates transcription re-initiation versus disassociation (Inlow et al. 2023; Brewer et al. 2025). Mutants in *rapA* have modest phenotypes including slow growth on medium containing high salt (Sukhodolets and Jin 1998), osmotic sensitivity (Fleurier et al. 2022) and biofilm defects (Lynch et al. 2007).

In this study, we examine *rapA* induction further using promoter/gene fusions and show that its inducibility by CPX is indeed independent of the LexA/RecA-dependent SOS response, and that its expression is repressed by DnaA; a protein that is both the replication initiation factor and transcriptional regulator. We examine the impact of RapA on survival to CPX, a topoisomerase II poison, in combination with SspA, another transcription factor implicated in the response to replication difficulties (Cooper et al. 2021; Sass et al. 2022; Sass and Lovett 2024). Our results show that these factors have opposing action: RapA promotes growth whereas SspA inhibits growth upon CPX treatment.When RNAse HI is inactivated, RapA promotes the ability to use RNA/DNA hybrids to initiate DNA replication whereas SspA interferes with this mechanism. SspA function appears to preclude or antagonize RapA’s positive effect on growth during CPX exposure and in the absence of SspA, RapA is no longer required for replication initiation at R-loops. We present a model where these proteins modulate the ability to clear elongation or paused complexes after replisome/RNAP collisions.

## RESULTS

### RapA expression is inducible by ciprofloxacin, independent of the SOS response and is under DnaA regulation

RNAseq analysis after replication inhibition via treatment with the chain terminating nucleoside, azidothymidine (AZT), showed a 4-fold elevation in expression after 1 hour of treatment (Sass and Lovett 2024); recent unpublished experiments with ciprofloxacin (CPX) showed a 26-fold enhancement after treatment of 50 ng/ml CPX. Induction was independent of LexA repressor cleavage, that underlies the well-studied SOS response to DNA damage (Simmons et al. 2008). Induction of *rapA* was noted previously for UV (Courcelle et al. 2001) and for norfloxacin in microarray studies (Dwyer et al. 2007). To examine the genetic regulation of *rapA* transcription, we fused its upstream promoter region to the *Photorhabdus* luciferase *luxCDEAB* operon (Van Dyk et al. 2001) so expression can be easily monitored by light production during culture growth. Strains were growth in rich medium and maintained in early exponential phase of growth, then challenged with CPX, with optical density (OD600) and luminescence monitored over time after treatment. We confirmed that expression from the *rapA* promoter was induced by CPX treatment in wild-type strains (Figure 1A). In *lexA3* mutant strains, in which the SOS response cannot be induced due to mutation at the cleavage site of the LexA repressor protein, or in RecA null mutants, induction was even stronger, confirming that RapA DNA damage induction is independent of the LexA/RecA-dependent SOS response. Hyper-induction in the *lexA3* background is seen for other SOS-independent genes such as the RpoS regulatory antiadaptor gene, *iraD*, (Sass et al. 2022), presumably because of longer persistence of damage in this repair-deficient strain background. Like *iraD*, induction of the *rapA* promoter was most pronounced after cultures approached saturation and stationary phase, suggesting some additional nutritional control of expression. Neither *lexA3* nor *sspA*Δ mutants altered constitutive expression, but both elevated expression after CPX treatment.

**Figure 1:**
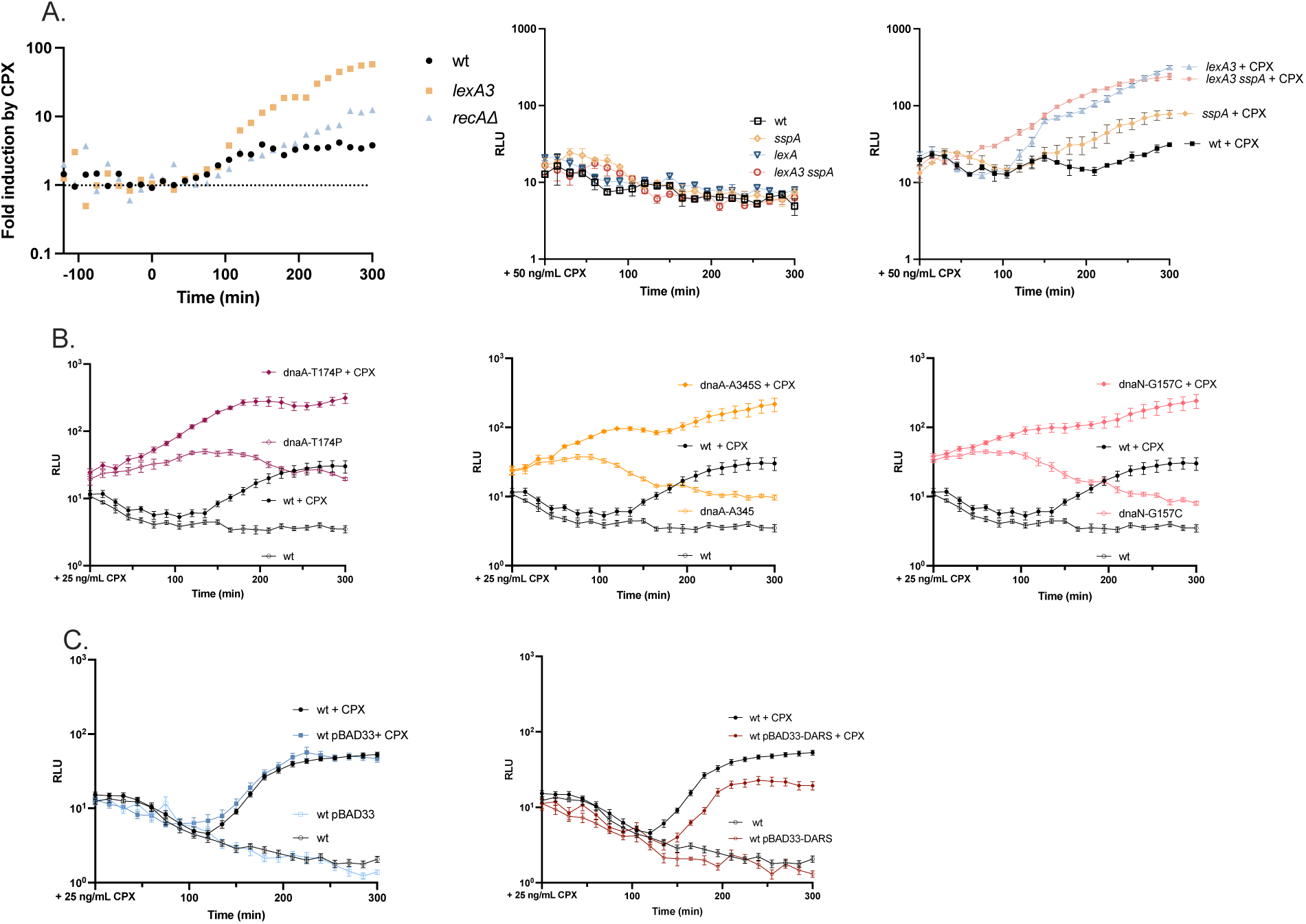
Luciferase expression from *rapA* promoter fusions. A. Induction of the *rapA* promoter by ciprofloxacin, as assayed by luciferase fusions. Ciprofloxacin (CPX) at 50 ng/mL was added at time 0 to early exponential phase cultures of the indicated genotype in LB. Relative luminesence units (RLU) were calculated by dividing luminescence by OD600 for each culture at each time point; fold induction by CPX was derived by dividing the average CPX-treated RLU by the average untreated RLU for each strain at each time point. B. As above, except 25 ng/mL CPX was used. In the *dnaA* and *dnaN* strains that promote a DnaA-ADP cellular environment, *dnaA* T174P, *dnaA* A345S, and *dnaN* G157C, the basal expression and inducibility of *rapA* is enhanced. C.When high copy of DARS1 sequence on vector pBAD33 is provided, yielding a DnaA-ATP cellular environment, *rapA* expression is repressed relative to vector alone.

Several SOS-independent DNA damage inducible promoters such as *iraD* and *nrdAB* (ribonucleotide reductase genes) are repressed by the DnaA protein in its ATP-bound state, with repression relieved through regulatory inactivation of DnaA (RIDA). RIDA is mediated through its interaction with the Hda protein and the replication processivity clamp (β, DnaN) (Gon et al. 2006; Sass et al. 2022). DnaA is the replication initiation protein of eubacteria, but also can act as a transcription factor (Messer and Weigel 2003). DnaA as a transcription factor is unusual in that it binds as a polymer, with high affinity binding promoted by its ATP-bound state (Erzberger et al. 2006).We tested whether expression of RapA was under similar genetic control by examining mutants that affect the nucleotide state of DnaA (Katayama et al. 2017). Three different mutants in *dnaA* and *dnaN* that promote the ADP-bound state of DnaA by stimulating its ATP hydrolyase activity led to markedly higher constitutive expression in luciferase fusion *rapA* promoter assays, consistent with DnaA-ATP repression. The *dnaA-*T174P and *dnaA-*A345S are constitutively inactivated and promote a cellular environment with DnaA in the ADP-bound state.We see *rapA* inducibility in these backgrounds and a higher level of basal expression. Similarly, the *dnaN-*G157C background promotes the DnaA-ADP form by favoring the Hda/ DnaN interaction which facilitates DnaA-ATP hydrolysis (Figure 1B).

Certain DNA sequences, known as “DnaA reactivation sequences” (DARS), can stimulate the DnaA exchange of ADP for ATP when present in higher copy; therefore, it would be expected to promote enhanced repression by DnaA-ATP (Fujimitsu et al. 2009). Indeed, *rapA* promoter-driven luciferase expression was lowered and/or induction suppressed by plasmids carrying the DARS1 locus, relative to vector only (Figure 1C). All together, these data indicate that inducibility of *rapA*, like that of *iraD* and *nrdAB,* is under DnaA control.

### RapA, SspA and growth in the presence of CPX

Given its strong inducibilty by CPX, we asked whether loss of *rapA*, either singly or in combination with *sspA* affected survival and proliferation after chronic exposure to CPX at three concentrations. In addition, because RNA/DNA hybrids are implicated in toxicity of replication/transcription conflicts (Aguilera and Garcia-Muse 2012; Gowrishankar et al. 2013; Hamperl and Cimprich 2014), we examined each genotype in combination with a deletion of *rnhA*, the gene which encodes RNase HI, an enzyme that removes RNA from RNA/ DNA hybrids (Figure 2A).

**Figure 2:**
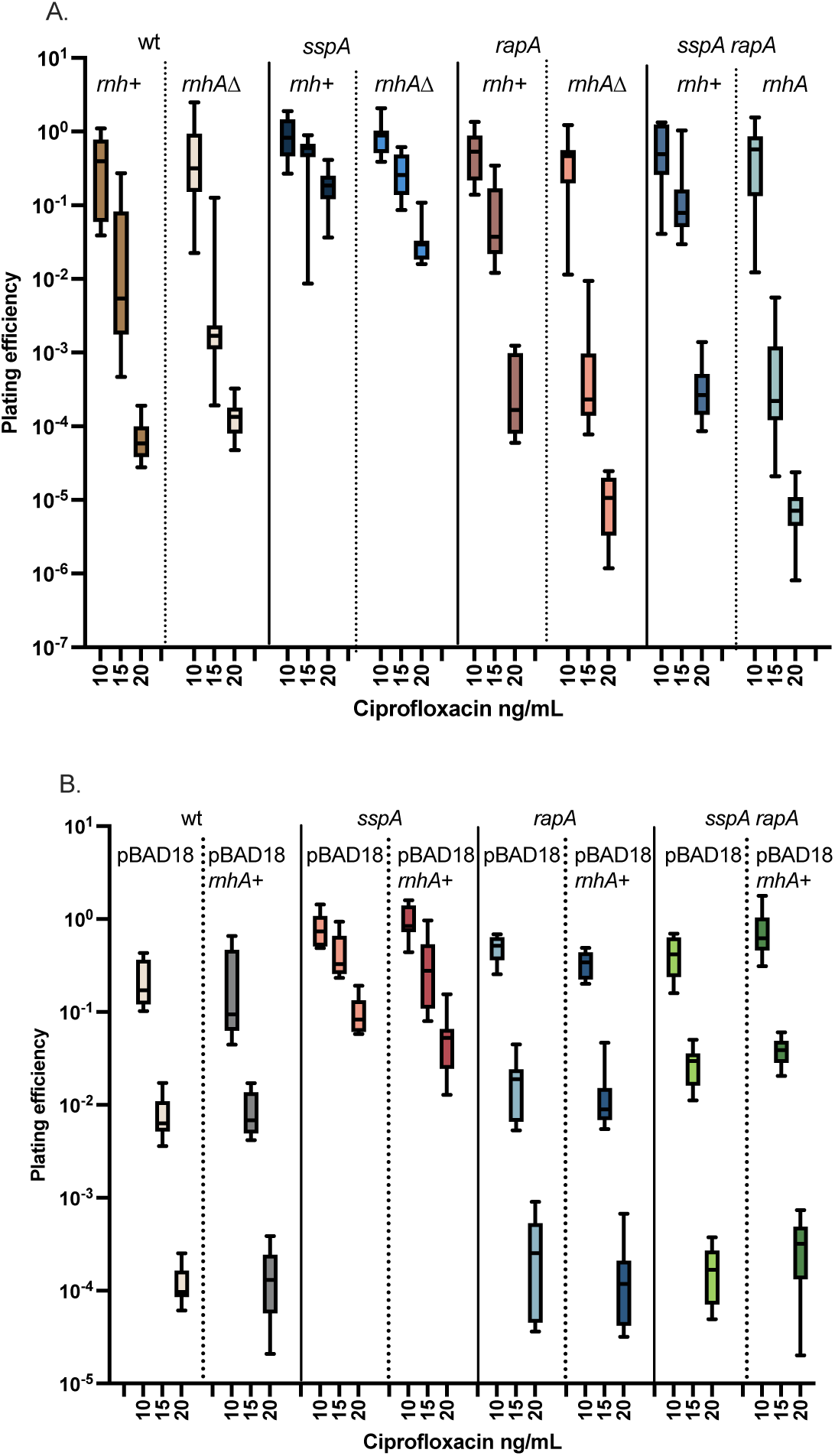
Plating efficiency of wt and mutant strains on LB medium containing ciprofloxacin at the indicated concentration relative to that on LB alone. Box plots indicate inter quartile distance; bars correspond median, minimum/maximum values for the number of independent cultures indicated. A. Strains with functional RNase HI, *rnhA*^+^, or lacking it, *rnhA*Δ, n = 10-17. B. Strains expressing RNase HI after arabinose induction from a plasmid, pBAD18-*rnhA*, or pBAD18 vector control, n = 8.

Mutants in *sspAΔ* were more resistant to CPX than wild-type strains suggesting that some SspA function is deleterious or inhibits growth under conditions of chronic replication stress. This is consistent with prior work showing that loss of SspA dramatically improves the poor viability of *holC* and *holD* mutants of the DNA polymerase III clamp loader complex (Michel and Sinha 2017; Cooper et al. 2021), another condition of chronic replication stress. Loss of *rapA* by itself did not significantly affect plating efficiency in the presence of CPX compared to *wt* strains. However, in the absence of *rnhA*, *rapAΔ* mutants showed increased sensitivity, over 10-fold relative to *rapA+ rnhAΔ* mutants at the higher CPX doses. Loss of *rnhA* did not sensitize *wt* strains to CPX; its effect was apparent only in *rapAΔ* strains. In addition, we found that the hyper-resistance of *sspAΔ* mutants to CPX was strongly dependent on RapA, suggesting an antagonistic relationship between SspA and RapA (Figure 3A). Overexpression of RNAse H1 did not significantly rescue the sensitivity of the *sspAΔ rapAΔ* mutant to CPX, implying that the role of RapA in promoting tolerance in *sspAΔ* mutants may be independent of R-loop avoidance (Figure 2B). The much larger effect of RapA in the *sspAΔ* mutant background suggests that SspA function interferes with that of RapA for CPX tolerance. We find a similar relationship between the effects of SspA and RapA on plating in the presence of replication inhibitor, AZT, (Fig. S1) confirming a general effect of the proteins on tolerance of replication stress.

**Figure 3:**
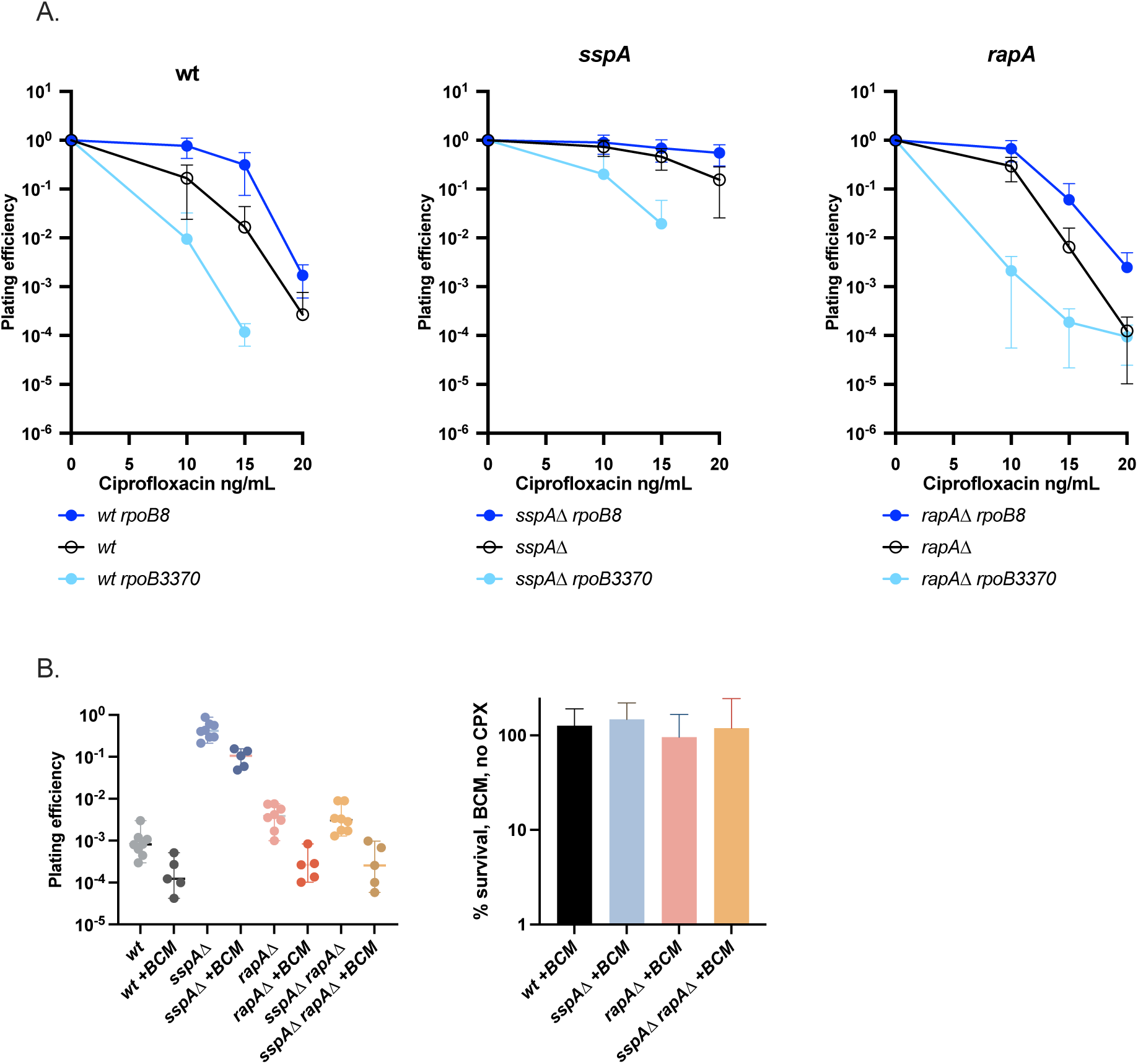
Ciprofloxacin (CPX) plating efficiency of strains carrying *rpoB* alleles affecting termination or with Rho inhibition by bicyclomycin. A. Effects of *rpoB* alleles. Error bars reflect standard deviations n = 8-14 B. Effects of bicyclomycin (BCM). Left, plating efficiency of the indicated strains on medium containing 15 ng/mL CPX ± 25 ng/mL BCM. Right, in the absence of CPX, % colony formation on 25 ng/ mL BCM vs no BCM. Bars indicate median, with 95% confidence intervals.

To implicate transcription properties in the phenotypes exhibited by mutants in transcription factors SspA and RapA, we combined mutants in these genes with alleles of the β subunit of RNA polymerase, *rpoB*, that alters elongation and termination properties (Figure 3A)(Jin et al. 1988; Jin and Gross 1991). The *rpoB8* allele, which increases pausing and termination propensity of RNAP, conferred additional resistance to CPX, whereas the *rpoB3370* allele, which increases processivity and diminishes termination propensity, conferred CPX sensitivity. Loss of *sspA* improves survival of both *rpoB* variants, whereas *rapAΔ* has little effect. The ability to clear TECs through termination is therefore important for tolerance of CPX lesions on DNA.

Prior work by Gottesman and colleagues has established that Rho-dependent termination is essential for survival after exposure to nalidixic acid (Washburn and Gottesman 2011), a quinolone compound with similar actions to CPX used here (Drlica and Zhao 1997). We used sublethal concentrations of the Rho inhibitor, bicyclomycin (BCM) to confirm, that Rho termination likewise contributes to tolerance of CPX treatment (Figure 3B). All genotypes examined, wt, *sspAΔ, rapAΔ,* and *sspAΔ rapAΔ* were sensitized to CPX by BCM, confirming the importance specifically of Rho-dependent termination in surviving replication stress.

Conflicts between the replisome and TECs are more deleterious in the head-on orientation, rather than when their movements along DNA are co-directional. To increase head-on collisions, the highly transcribed *rrnA* operon was inverted, creating the “InvA” strain (Boubakri et al. 2010); the strain is normally viable except in the absence of helicases that are implicated in TEC removal. Inversion of the *rrnA* operon produces significantly more sensitivity to chronic CPX exposure compared to wt, sensitivity that is entirely reversed by loss of *sspAΔ.* (Figure 4A). InvA also sensitizes *rapAΔ* mutants, but has little effect on *sspAΔ* mutants. Loss of *rapA* produces pronounced sensitivity when combined with *sspAΔ* background but, interestingly, InvA has little additional effect in this background, suggesting that inviability promoted by the *rrnA* inversion requires SspA. This indicates that with increased head-on collisions, RapA’s RNAP role is beneficial, whereas that of SspA is deleterious, with SspA antagonizing RapA’s positive effects. Overexpression of RNase HI in strains carrying the inverted *rrnA* orientation fails to rescue CPX sensitivity conferred by InvA (Figure 4B), showing that it is not due to increased R-loop formation; in fact, RNase H sensitizes *sspAΔ* InvA strains (p < 0.01, Mann Whitney test), showing that RNA/DNA hybrids in some way promote tolerance of the inversion, in a fashion dependent on RapA.

**Figure 4:**
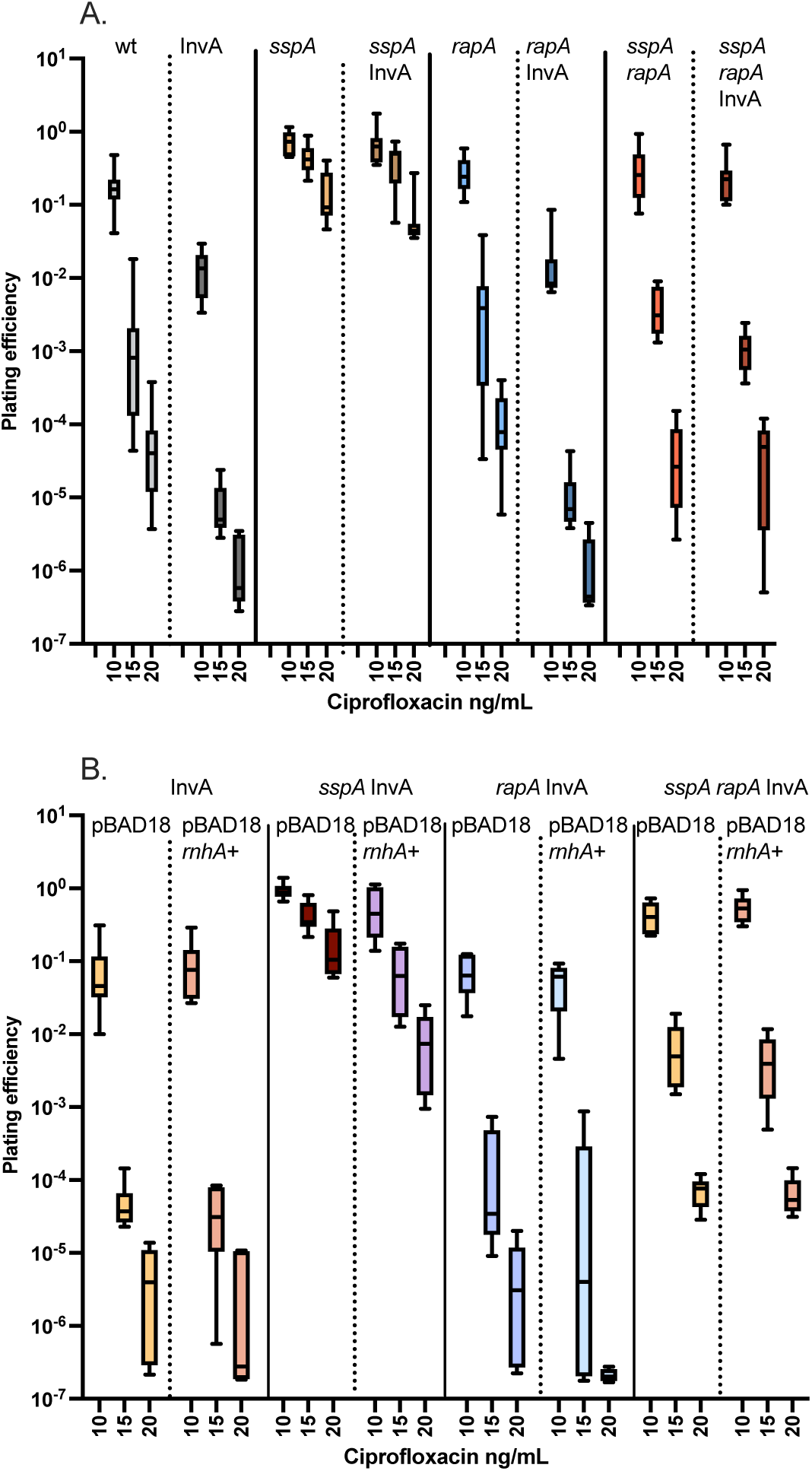
Ciprofloxacin (CPX) plating efficiency in strains carrying an inversion of *rrnA* region, InvA. Box plots indicate median and bars correspond minimum/maximum values. A. Comparison between wt and InvA inverted strains, n = 8-16. B. InvA strains with arabinose-induced expression of RNase HI from pBAD18 *rnhA*+ compared to pBAD18 vector alone, n = 6.

The synergistic effect of *rnhAΔ* and *rapAΔ* on CPX tolerance suggests that RapA may inhibit R-loop formation, something suggested from recent biochemical analysis of RapA with RNAP (Brewer et al. 2025). To examine this further in vivo, we took advantage of observations by Kogoma and colleagues, that R-loop formation in *rnhAΔ* mutants can drive initiation of replication, independent of the initiator protein, DnaA, and the origin of replication, *oriC* (Kogoma 1997). We employed a temperature-sensitive allele of DnaA, *dnaA46*, that reduces plating efficiency on LB at 39° versus 30° over 4 orders of magnitude (Figure 5). When *rnhAΔ* mutants are introduced into this *dnaA46* background, plating efficiency at 39° is almost normal, due to efficient R-loop initiation of replication in the absence of degradation by RNase HI. Mutants in *sspAΔ* elevated suppression of *dnaA46* by *rnhAΔ* even further, whereas loss of *rapA diminished* suppression. Rather than acting merely to prevent R-loop formation as expected from the biochemical analysis, RapA function is required to utilize R-loops for replication, possibly because it promotes RNAP polymerase dissociation from TECs while leaving potential RNA primers for DNA synthesis behind. The double *sspAΔ rapAΔ* mutant resembles *sspAΔ* in promoting R-loop initiation; that is, RapA function in promoting R-loop initiation is only required in the presence of functional SspA. Interestingly, *sspAΔ* mutants have some ability to suppress *dnaA46* at 39° even in the presence of functional RNase HI, although this property is variable among the populations and could represent fast-arising genetic or epigenetic variants. These observations are consistent with the hypothesis that SspA promotes stable (stalled or terminated) RNAP complexes that impede replication, with nascent RNA inaccessible for priming DNA synthesis. RapA can promote release of these complexes, lending access to RNA for initiation. When SspA is absent, release must frequent enough without RapA function.

**Figure 5:**
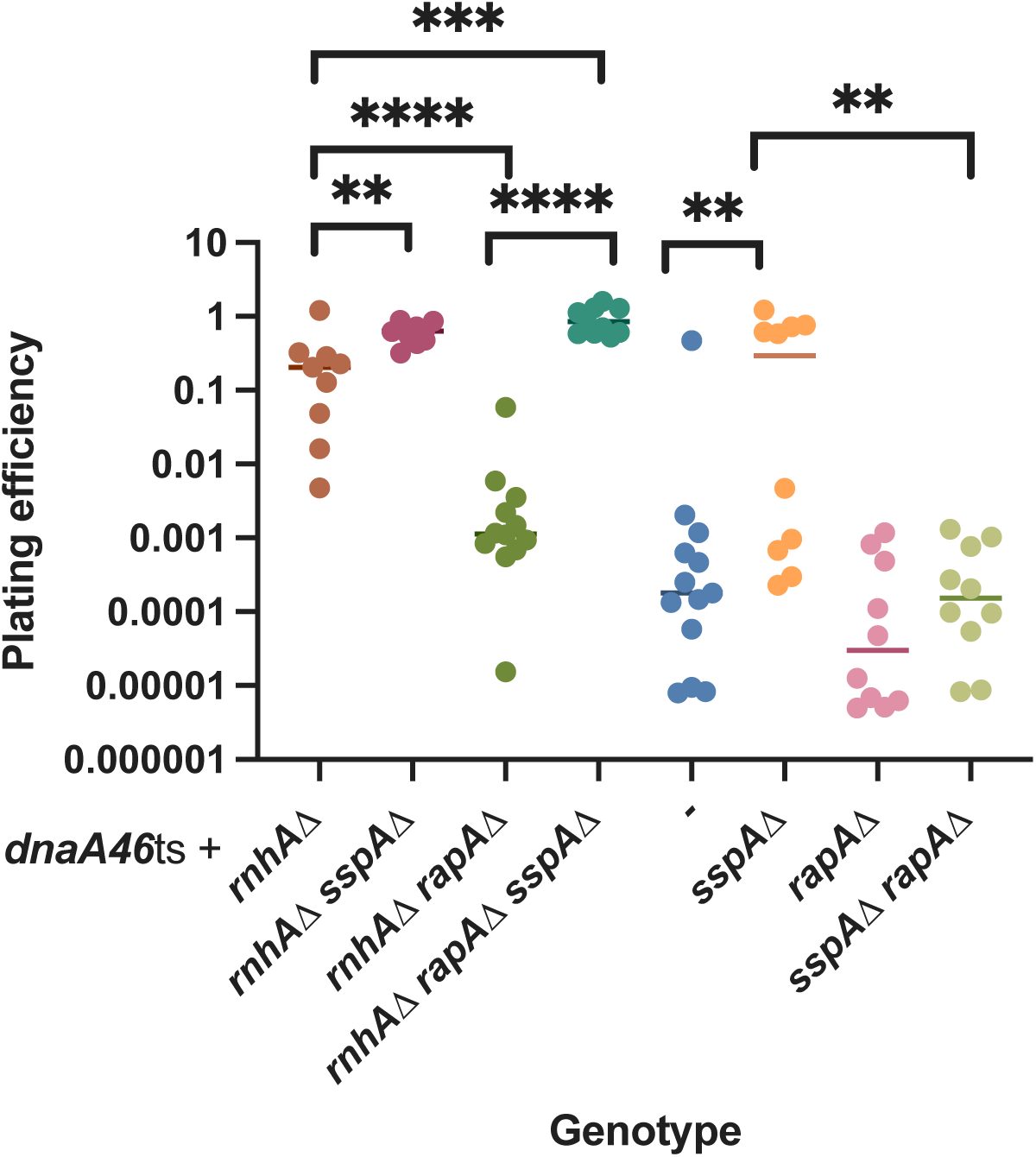
The ability to use R-loops to initiate replication independent of DnaA. Shown are plating efficiencies at 39° relative to 30° for strains carrying *dnaA46* and the additional genetic alleles indicated. Significance values indicate derive from pairwise Mann Whitney tests: ** with p < 0.01; *** with p < 0.0001; **** with p < 0.00001.

## DISCUSSION

DNA damage, such as that after CPX treatment, leads to the accumulation of single-strand DNA (ssDNA) post-replication gaps due to “lesion-skipping” by the replisome (Cox et al. 2023). Ongoing transcription on such templates would be problematic: superhelicity is lost and RNAP may be trapped by encountering strand breaks on the template strand. Strand interruptions on the non-template strand can also lead to loss of the displaced DNA strand behind transcription elongation complexes (TECs), allowing extensive RNA/DNA hybrid formation due to the loss of the competitor strand for pairing.

Clearing these trapped complexes may be necessary to fill daughter-strand gaps to complete replication and stabilize the chromosome.

This study shows that inability to clear TECs caused by mutations in RNAP or inhibition of Rho-dependent transcription termination correlates with compounded sensitivity to DNA damaging agents such as the topoisomerase II inhibitor, ciprofloxacin (CPX). Inversion of a highly transcribed rRNA operon such that the replisome and TEC collide head-on, also impairs CPX tolerance.This work implicates two RNA-polymerase associated proteins, RapA and SspA in regulating clearance, with RapA playing a positive and SspA a negative role, often in opposition to one another.

Both RapA and SspA are under cell growth regulatory control, in opposite fashions. RapA had previously been shown to be growth-rate and growth-phase regulated, with highest expression in early exponential phase of growth (Cabrera and Jin 2001). We show here that it is under repression by DnaA, and is induced upon treatment with replication inhibitors, such as CPX. SspA, on the other hand, is positively regulated by the stringent response to starvation, induced by accumulation of ppGpp in late exponential phase of growth (Williams et al. 1994), although it is abundant even at earlier phases (Dolgalev et al. 2023). In their opposing effects on replication stress tolerance, RapA likely promotes a state conducive to cell proliferation (consistent with its early log prevalence) whereas SspA promotes an anti-proliferative state (consistent with its induction by starvation). Protein abundance studies show that SspA concentrations predominate over those for RapA by about 10-fold (Dolgalev et al. 2023), but that value may be highly sensitive to growth conditions.

RapA protein is a SWI2/SNF2 protein that was found to co-purify with RNAP and to promote polymerase recycling, dependent on ATP hydrolysis, during in vitro transcription reactions (Muzzin et al. 1998; Sukhodolets and Jin 2000; Sukhodolets et al. 2001). Unlike many SWI/SNF proteins it only poorly binds DNA, with its ATPase stimulated upon RNAP interaction (Sukhodolets and Jin 2000; Sukhodolets et al. 2001). It binds preferentially to core RNAP and is readily displaced by σ70 (Shaw et al. 2008).

Single-molecule studies show that it binds RNAP that is sequestered in a post-termination complex after RNA release and accelerates its dissociation from DNA, by promoting opening of the β-β’ clamp (Inlow et al. 2023). In that study, RapA did not bind appreciably to elongation complexes; structural studies suggest that the presence of RNA in the exit channel of RNAP should impede RapA binding (Inlow et al. 2023; Brewer et al. 2025). On negatively supercoiled DNA, PTCs can initiate σ-independent non-specific transcripts, which are more prone to R-loop formation and intrinsic terminator read-through (Brewer et al. 2025). Consistent with the idea that RapA prevents toxic R-loop formation in vivo by destabilization of PTCs, *rapAΔ* mutants are modestly sensitive to the Rho-termination inhibitor bicyclomycin, with rescue by overproduction of RNase HI; the salt-sensitive phenotype of *rapAΔ* mutants is also relieved by RNase HI overexpression (Brewer et al. 2025).

Our finding reported here that RapA promotes tolerance to CPX in *rnhA* mutants, is consistent with a role in preventing some form of toxic R-loop formation.This simple hypothesis, however, does not explain other aspects of our results. RapA is strongly required for CPX tolerance in *sspA*Δ mutants, which is not relieved by RNase HI overexpression, suggesting that RapA plays a role independent of R-loop formation. Furthermore, RapA *promotes*, in some way, the creation or use of R-loops to initiate DNA replication independent of DnaA. Elongation factor DksA also is required for R-loop initiated replication (Myka et al. 2019); based on genetic interactions in that study, DksA has been proposed to destabilize stalled complexes, releasing RNAP but not the RNA/DNA hybrid. One explanation for our results may be that RapA can also promote release of RNAP from certain stalled TECs, in this case leaving behind a RNA/DNA hybrid molecule, possibly one that has backtracked or that has lost its displaced DNA strand because of transcription on a nicked or gapped chromosome. Release of stalled RNAP by RapA through backtracking has been proposed based on one study (Portman et al. 2022).

SspA was discovered as a highly abundant protein associated with RNA polymerase of *E. coli* (Ishihama and Saitoh 1979) and is found ubiquitously across eubacterial species (Wang et al. 2021). SspA associates preferentially with the RNAP σ^70^-holoenzyme over the core enzyme (Ishihama and Saitoh 1979) or σ^s^-holoenzyme (Wang et al. 2020). In vivo, SspA can act as a transcriptional activator, for example at the *gadA* acid response gene promoter, or as a repressor, for example at peptidoglycan synthase genes, *ponA* and *rodA* (Lou et al. 2022). Structural studies show that *E. coli* RNAP SspA does not directly contact DNA but forms tight interactions with σ70. This ability to stabilize σ70 complexes explains how SspA acts as an activator for promoters with relatively poor-35 sequences such as *gadA* (Travis et al. 2021), but could act as a repressor for other promoters by impeding promoter clearance during initiation for others (Wang et al. 2020). Although most work has concentrated on the role of SspA in initiation, it likely plays additional roles as well, particularly because it interacts with the β’ zinc binding domain (ZBD), a region of RNAP known to modulate termination properties (Hu and Liu 2022), and with ω (Wang et al. 2020;Travis et al. 2021). Its genetic interactions with RapA, which associates strictly with core RNAP (Shaw et al. 2008), and mutants of β that influence elongation and termination properties, suggests the effects we see of SspA on CPX plating efficiency and R-loop initiated DNA replication derive from SspA effects on elongation or termination rather than initiation.

Structural information for RapA/RNAP and SspA/RNAP complexes show that both RapA and SspA bind to a similar region, near the ZBD of β’ protein; therefore their occupancy in RNAP is expected to be mutually exclusive. In this case, paused TEC or post-termination complexes (PTCs) carrying SspA should be refractory to release by RapA, explaining why we see little contribution to CPX survival unless SspA is deleted. Inverting a single rRNA operon, *rrnA*, was sufficient to reduce CPX tolerance when SspA is functional; this could be due to the persistence of PTCs at this site. Interestingly this inversion is associated with a pronounced replication stall site near the intrinsic terminator of the *rrnA* operon when the Rep helicase is absent (Boubakri et al. 2010). Rep may normally be clearing SspA-containing PTCs but fail to be recruited to the *rrnA* inversion site after lesion skipping induced by CPX, because it remains associated with DnaB at the fork (Guy et al. 2009) that has moved on.

Although SspA appears to be deleterious for proliferation under conditions of chronic replication stress, a role in stabilizing stalled TECs on DNA could help survive impending or ongoing starvation, allowing cells to complete transcription that would otherwise abort, before entering quiescence. Indeed, SspA is required for long-term survival after starvation (Williams et al. 1994). In addition, a role in stabilizing TEC against release by RapA may allow more efficient re-initiation at certain loci, in a strictly σ70-dependent fashion because of its preferential association with σ70-holoenzyme. In *Francisella tularensis*, SspA is recruited through interactions with co-activator proteins (Cuthbert et al. 2017) and in *E. coli,* SspA is co-activated by interaction with bacteriophage P1 Lpa protein (Hansen et al. 2003).Whether *E. coli* possesses other co-activators for SspA remains to be determined. The loading of SspA at specific initiation sites may constitute a locus-specific mechanism for controlling elongation and termination properties in a locally-restricted way.

Other RNAP-associated factors that modulate elongation and termination properties also contribute to survival of replication stress.The elongation factor DksA and the alarmone ppGpp are required for optimal resistance to UV light, nalidixic acid and other DNA-damaging treatments (McGlynn and Lloyd 2000; Meddows et al. 2004; Sivaramakrishnan et al. 2017; Myka et al. 2019) and many others remain to be investigated.

## MATERIALS AND METHODS

### *E. coli* K-12 strains and growth

All strains used are MG1655 and mutants were constructed via P1 transduction (Miller 1992), selecting for a linked antibiotic resistance marker. Appropriate antibiotic concentrations used were kanamycin (Kn, 20 μg/ml), chloramphenicol (Cm, 15 μg/ml), ampicillin (Ap, 100 μg/ml), and/or tetracycline (Tc, 10 μg/ml).The LB Lenox formulation (Miller 1992) was used for standard growth media with the addition of 2% Bacto-agar for plate formation.

### Plasmid construction

Plasmid pBAD33-DARS1 was constructed by digesting the pBAD33 vector (Addgene #36267) with HindIII and SacII restriction enzymes.The DARS1 insert was created by annealing oligo DARS1_1 (5’- agctt gggat agggg ctgga gacag ttatc cacta ttcct gtgga taacc atgtg tatta gagtt agaaa acacg agct - 3’) and DARS1_2 (5’ - cgtgt tttct aactc taata cacat ggtta tccac aggaa tagtg gataa ctgtc tccag cccct atccc a - 3’) then ligating the plasmid backbone and insert with T4 ligation (New England Biolabs) before transforming into competent *E. coli.* The *rapA* promoter luciferase plasmid was constructed by Gateway Cloning (Invitrogen) using the primer pair (5’ - ggggA CAAGT TTGTA CAAAA AAGCA GGCTT CCTCG CCTTG CCGAT GAAGA AAACC AAAAG CGTGG T - 3’) and (5’ - ggggA CCACT TTGTA CAAGA AAGCT GGGTC CTAAT GTGTT CGGCT CTATA TCTTT AATTG CAGGC AAT - 3’) to amplify the *rapA* promoter region with Phusion (New England Biolabs).The Gateway BP reaction inserted the fragment into the pDONR221 vector and the LR reaction moved the *rapA* promoter into the pDEW201vector (Van Dyk et al. 2001) as previously described (Merrikh et al. 2009). The pBAD18-rnhA plasmid was created by amplifying *rnhA* (5’ - GAGGC TAGCA TGCTT AAACA GGTAG AAATT TTCAC C - 3’) and (5’ - GGCAA GCTTT TAAAC TTCAA CTTGG TAGCC - 3’) using Phusion.The fragment and vector were digested with NheI and HinDIII (New England Biolabs) according to the manufacture’s protocol and ligated with T4 ligase before transforming into competent *E. coli*.

### Luciferase expression assays

Luminescence and OD600 were measured using BioTek Cytation 1 Plate Reader and Costar 96 Well Assay Plate (treated polystyrene, black plate, clear bottom) as previously described (Sass et al. 2022). Briefly, colonies were grown to log phase shaking in LB media with appropriate antibiotics. Cultures were then inoculated in the 9- well plate at 1:100 dilution and grown for 2 hours prior to treatment with 25 ng/mL ciprofloxacin. Relative luminescence units (RLU) was calculated by normalizing the bioluminescence to the OD600 every 15 minutes. Data are averages of independent replicate cultures.

### Plating efficiency assays

Overnight cultures were normalized in LB media and serially diluted 10-fold in 1×56/2 salts before plating 5 μL on the assay plates and incubated for approximately 16 hours at 37°C unless otherwise specified for temperature-dependent assays. Colony forming units (CFU) were quantified to determine the plating efficiency at each condition when normalized to the no-drug control or permissive temperature plate.The indicated ciprofloxacin (CPX) concentration was added the LB assay plates containing 2% (wt/vol) agar.To induce *rnhA* over expression in the pBAD18 plasmid vector, plates were supplemented with 0.2% arabinose and appropriate antibiotics. 25ng of bicyclomycin (BCM) was spread evenly on the assay plates and dried prior to plating the serially diluted cultures.

## ACKNOWLEDGMENTS

This word was supported by NIH grant R01 GM51753 to STL and T32 007122 to THS.. FIGURE LEGENDS

---

**TABLE 1.**
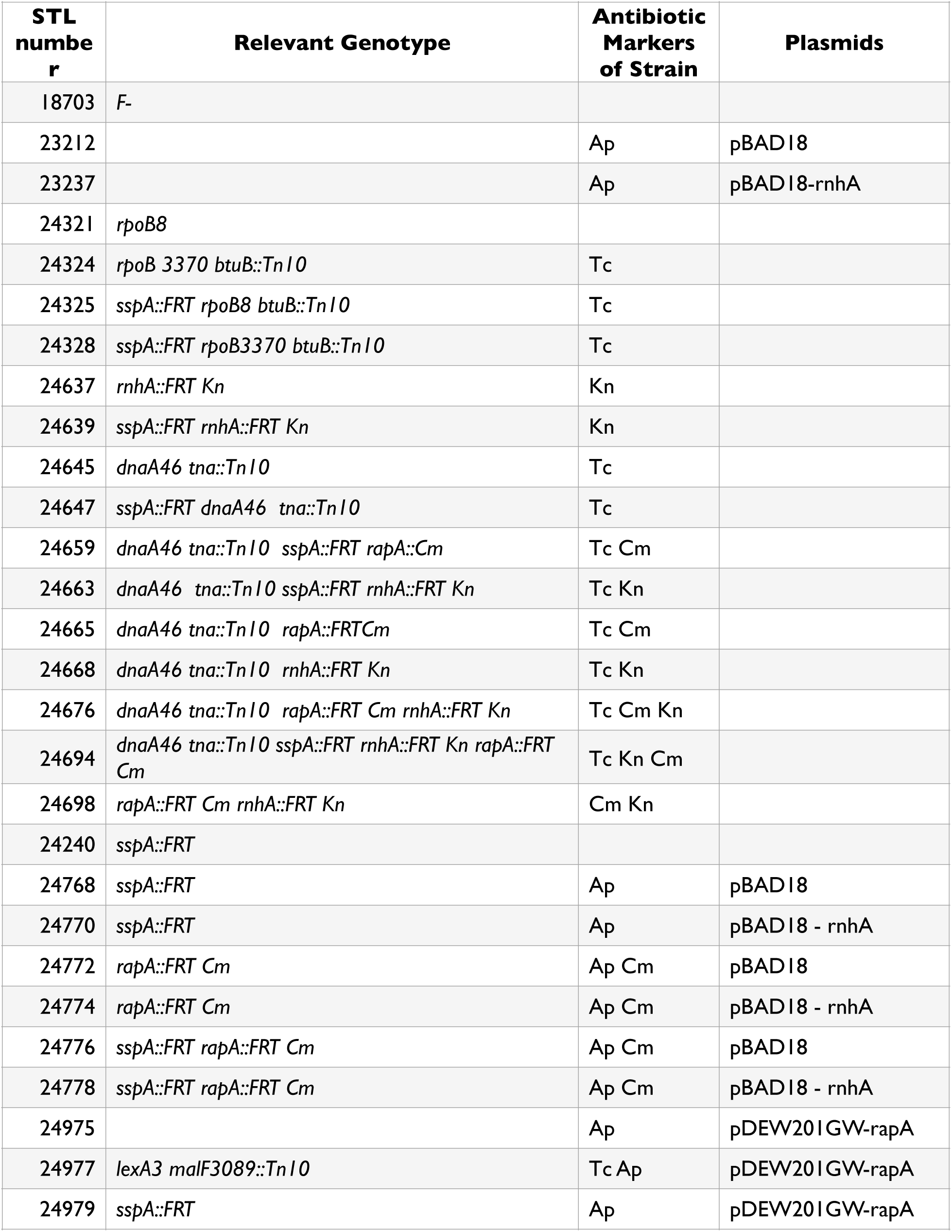
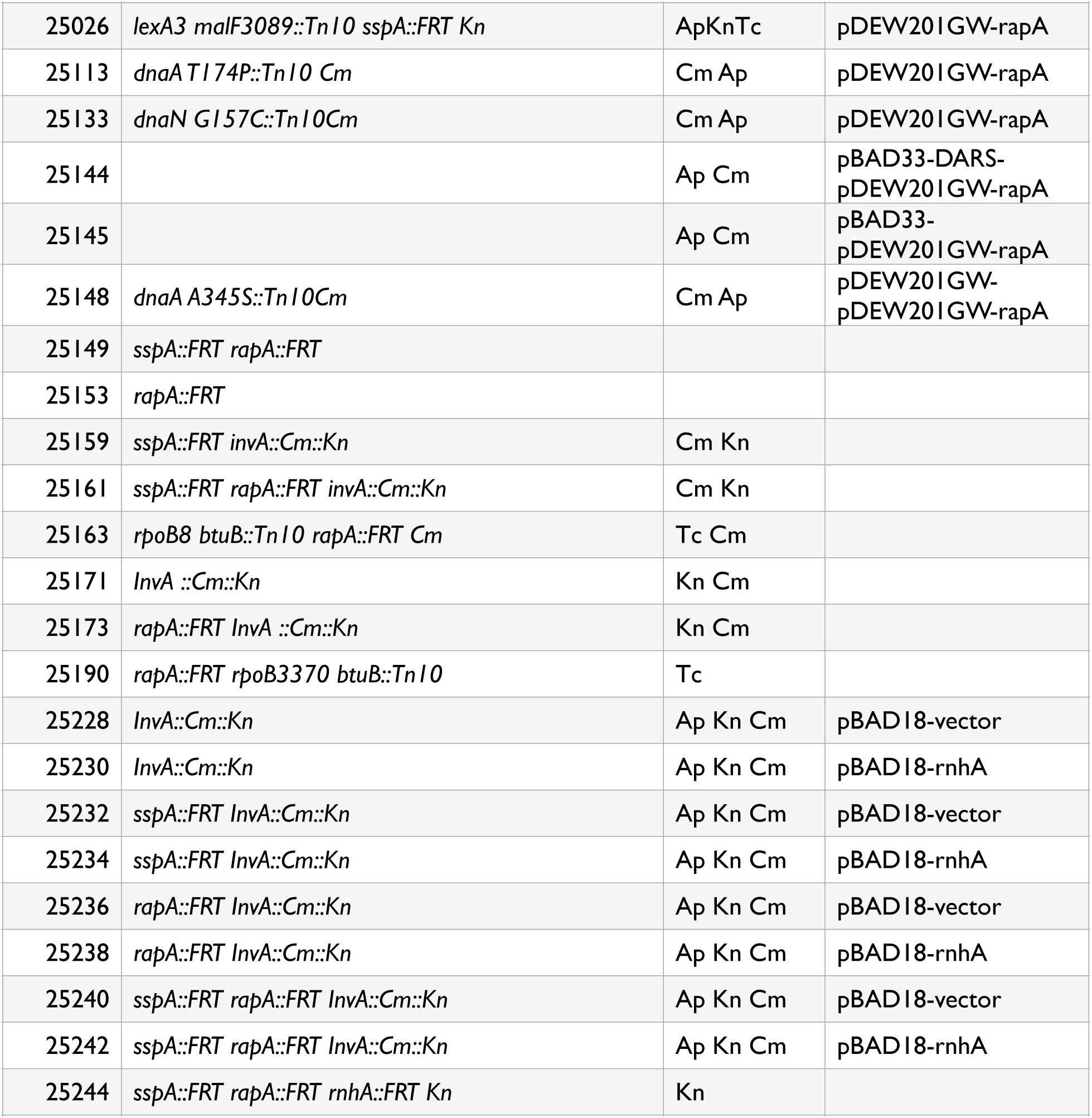
STRAIN LIST.

