## Supplementary figures and images for "*Escherichia coli* transcription factors RapA and SspA play opposing roles in tolerance to replication/ transcription conflicts after DNA damage"

### Supplemental Fig.1

Supplemental Figure 1.  Plating efficiency of strains on medium containing azidothymidine (AZT).

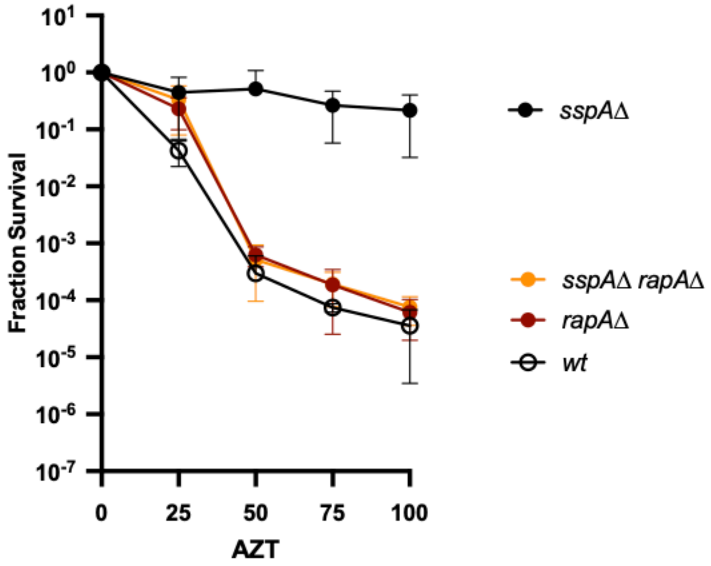
